# Utility of thromboelastography with platelet mapping (TEG-PM) for monitoring platelet transfusion in qualitative platelet disorders

**DOI:** 10.1101/2024.05.09.593198

**Authors:** Robert H Lee, Tanvi Rudran, Wolfgang Bergmeier

## Abstract

Patients with pathogenic variants in *RASGRP2* (inherited platelet disorder (IPD)-18) have normal platelet counts but show impaired platelet aggregation due to diminished activation of αIIbβ3 integrin. This defect results in moderate to severe bleeding episodes, especially following surgical procedures, which require patients to be transfused with platelets and/or pro-hemostatic agents. We recently demonstrated that the hemostatic efficacy of transfused platelets is limited by dysfunctional endogenous platelets in a mouse model of IPD-18 (*Rasgrp2*^*-/-*^ mice), as dysfunctional platelets were recruited to the forming hemostatic plug but did not participate in clot contraction. Consequently, higher amounts of transfused platelets were required to outcompete these dysfunctional cells and to reverse bleeding. We here studied the usefulness of thromboelastography with platelet mapping (TEG-PM), a method to evaluate platelet-dependent clot contraction, for *ex vivo* monitoring of the hemostatic potential in *Rasgrp2*^*-/-*^ mice transfused with various amounts of wild-type (WT) platelets. *Rasgrp2*^*-/-*^ whole blood samples did not contract in TEG-PM, consistent with a critical role of this protein in αIIbβ3 activation. Addition of WT platelets improved TEG parameters (K time, α-angle, MA) in a ratio dependent manner, consistent with our recent in vivo studies showing impaired hemostasis at a 5:1, but not at a 2:1 ratio of mutant to WT platelets. Interestingly, K and α values were identified as better predictors of transfusion efficacy than MA, the most platelet-dependent TEG parameter. In conclusion, this proof-of-concept study supports the use of TEG-PM to monitor platelet transfusion ratios and hemostatic potential in IPD-18 and potentially other platelet disorders.

## Introduction

Platelet transfusion therapy is instrumental in preventing major hemorrhage in patients with hypoproliferative thrombocytopenia [1]. One transfusion unit can increase the circulating platelet count by more than 30,000/µl, easily maintaining patients above a critical threshold of 10,000/µl [2]. In these patients, platelet recovery in circulation post-transfusion is often monitored by methods which approximate the increase in platelet count, such as corrective count increment (CCI) and percent platelet recovery (PPR) [3]. However, these methods have limited accuracy in patients with higher pre-transfusion platelet counts, such as patients with platelet function disorders (PFDs), and they do not determine the origin of the platelets measured (i.e., endogenous vs. transfused).

PFDs include acquired disorders such as from antiplatelet therapy, or inherited disorders such as Glanzmann’s thrombasthenia (GT) or inherited platelet disorder (IPD)-18. IPD-18 is caused by pathogenic variants of *RASGRP2*, the gene encoding for calcium- and diacylglycerol-regulated guanine nucleotide exchange factor I (CalDAG-GEFI), a protein which is highly expressed in platelets and plays a critical role in inside-out activation of integrin αIIbβ3 [4]. IPD-18 patients can suffer from severe spontaneous and surgical bleeding requiring pro-hemostatic therapies such as recombinant activated FVII (rFVIIa), anti-fibrinolytics and platelet transfusion [5,6]. Interestingly, several groups have reported an inability of platelet transfusions to control bleeding [6–8], potentially due to the low transfused to endogenous platelet ratio as has been suggested in GT patients [9,10]. We recently performed mechanistic studies in *Rasgrp2*^*-/-*^ mice, which demonstrated that endogenous, dysfunctional platelets in these mice impair the hemostatic activity of transfused, healthy platelets by two mechanisms: 1) competition for primary adhesion sites, and 2) impaired clot contraction. As a result, a 2:1 ratio of dysfunctional to healthy platelets was required for hemostatic success [11]. In that study, platelet ratios were monitored by flow cytometry, a technique that is not routinely used clinically especially when studying disorders of intracellular proteins such as CalDAG-GEFI. We here investigated whether thromboelastography with platelet mapping (TEG-PM), a method that evaluates platelet function in the context of clot contraction, can be used to monitor platelet ratios and hemostatic potential in *Rasgrp2*^*-/-*^ mice following platelet transfusion.

## Materials and Methods

### Mice

*Rasgrp2*^*-/-*^ [12] and *Talin1*^*fl/fl*^ *Pf4-Cre*^*+*^ (*Talin1*^*mKO*^) [13] were previously described and maintained in-house on a C57BL/6J background. Wild-type (WT) mice were C57BL/6J from Jackson Laboratory. Male and female mice between 8-16 weeks were used for experiments. All procedures were approved by the Animal Care and Use Committee of the University of North Carolina at Chapel Hill.

### Aggregometry

Citrated platelet rich plasma (PRP) from WT and *Rasgrp2*^*-/-*^ mice was prepared by diluting citrated whole blood (WB) with 1/3 volume of modified Tyrode’s buffer (in mmol/L; 137 NaCl, 0.3 Na_2_HPO_4_, 2 KCl, 12 NaHCO_3_, 5 HEPES, 5 glucose, pH 7.3) followed by centrifugation at 120 x g for 4 mins. PRP adjusted to 2.5 x 10^8^/ml with additional Tyrode’s buffer was mixed to achieve specific platelet ratios (or kept separate) and aggregated with ADP (10 µM, Sigma) or type I collagen (10 µg/ml, Chronolog).

### TEG with Platelet Mapping (TEG-PM)

Heparinized WB samples were collected from the retroorbital plexus using heparinized glass capillary tubes (Fisher) and were left to rest for 20 minutes prior to running the TEG-PM assay. Platelet counts in WB were determined by flow cytometry and WB volumes were mixed to achieve specific platelet ratios or kept separate. WB samples were then added to cups containing TEG-PM reagents (activator F (reptilase + FXIIIa) and arachidonic acid) and immediately analyzed with a TEG-5000 Hemostasis Analyzer (Haemonetics). Individual TEG parameters were recorded for each sample: K, α-angle and MA, which measure time to minimum clot strength, rate of clot formation/contraction and maximal clot strength, respectively.

### Flow cytometry platelet activation

Diluted heparinized WB samples (2 µl WB in a 25 µl reaction mixture) were incubated for 15 minutes with 20 µM ADP in the presence of anti-GPIX-AlexaFluor(AF)647 and JON/A-PE (Emfret Analytics, both conjugated in-house, 2 µg/ml final concentration), then diluted 1:5,000 in PBS and analyzed on a BD Accuri C6 Plus flow cytometer. ≥95% WT platelets and <5% *Rasgrp2*^*-/-*^ platelets become JON/A-PE positive upon ADP stimulation, so % WT platelets in the sample can be estimated, with some small amount of error.

### Confocal microscopy

In some samples prior to TEG-PM, anti-GPIX antibodies (5 µg/ml) and AlexaFluor647-labeled human fibrinogen (Life Technologies, 100 µg/ml) were added to WB. Formed clots were gently removed from TEG cups, fixed in 2% formaldehyde overnight, and Z stacks of slide-mounted clots were obtained by laser scanning confocal microscopy (Leica DM4000) using a 63X oil objective.

### Statistics

TEG-PM parameter data shown as mean ± standard deviation. Normality was determined by Shapiro-Wilk test. Parametric data was analyzed by one-way ANOVA with Tukey’s multiple comparisons test, and non-parametric data by Kruskal-Wallis with Dunn’s multiple comparisons test. P < 0.05 was considered statistically significant.

## Results and Discussion

We recently demonstrated that hemostasis is restored in *Rasgrp2*^*-/-*^ mice when post platelet transfusion ratios of endogenous (*Rasgrp2*^*-/-*^) to transfused (WT) platelets were in the range of 2:1, but not when in the range of 5:1 [11]. We first tested whether standard aggregometry could be used to monitor the hemostatic potential in these mice. 5:1 or 2:1 ratios of *Rasgrp2*^*-/-*^ to WT platelets in PRP were established in vitro and samples were stimulated with ADP or collagen, agonists that induce little to no aggregation response in *Rasgrp2*^*-/-*^ platelets (Figure 1A,B) [12]. Adding in WT platelets at either ratio only minimally improved aggregation (Figure 1A,B), as *Rasgrp2*^*-/-*^ platelets did not interact with WT platelets and remained in suspension, preventing light transmission (Figure 1C). Thus, aggregometry did not prove to be a useful method to monitor the hemostatic potential of healthy platelets in the presence of platelets with impaired integrin activation.

**Figure 1:**
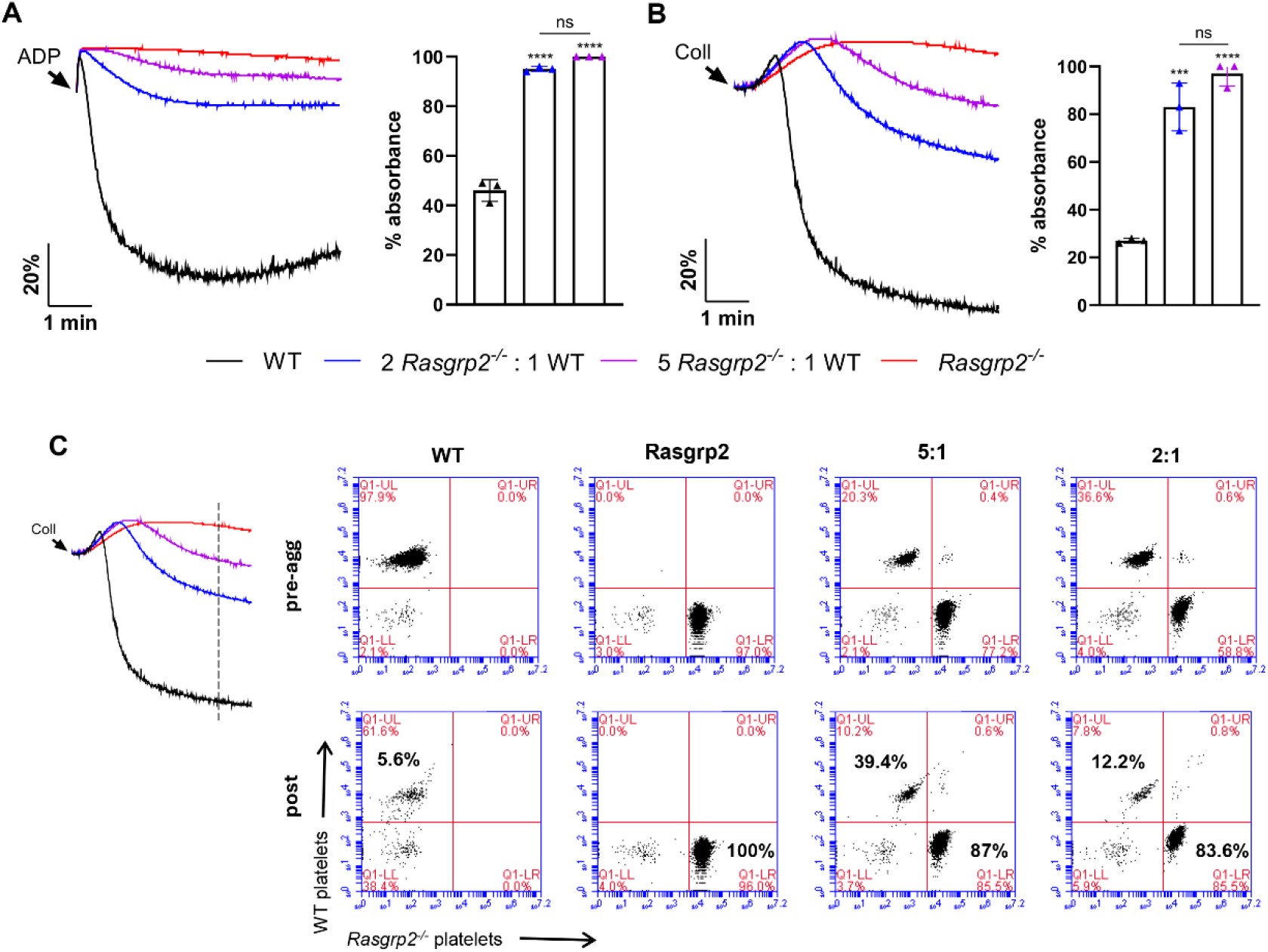
(A,B) Platelet aggregometry studies with mixed WT and *Rasgrp2*^*-/-*^ samples. Platelet rich plasma (PRP) was prepared from WT and *Rasgrp2*^*-/-*^ mice and analyzed by light transmission aggregometry separately or mixed to achieve specific platelet ratios (2:1 or 5:1 *Rasgrp2*^*-/-*^ to WT platelets). Platelets were aggregated with 10 µM ADP (A) or 10 µg/ml type I collagen (B), and peak light absorbance was determined for n=3 experiments (100% absorbance = no aggregation). Statistical significance was determined by one-way ANOVA with Tukey’s multiple comparisons. *** P<0.001, **** P<0.0001. (C) In some experiments prior to aggregation, WT and *Rasgpr2*^*-/-*^ PRP were differentially labeled with AlexaFluor conjugated anti-GPIX antibodies. Aliquots were removed from cuvettes prior to agonist addition and near peak aggregation (indicated by dotted line), and WT and *Rasgrp2*^*-/-*^ platelets in suspension were determined by flow cytometry (gated for single platelet population).

Our in vivo studies identified defective clot contraction as a major reason for impaired hemostasis in transfused *Rasgrp2*^*-/-*^ mice [11]. We also demonstrated that standard TEG is insensitive to defects in the Rap1 GTPase signaling pathway in platelets, including CalDAG-GEFI deficiency [14]. We therefore evaluated whether Platelet Mapping TEG (TEG-PM) could be used to monitor the hemostatic potential in these mice. Samples were activated with arachidonic acid and we analyzed K time, α-angle (α) and MA (see methods). Clot formation was severely impaired in whole blood from *Rasgrp2*^*-/-*^ mice when compared to controls, with MA values <10 suggesting a near-complete absence of platelet function (Figure 2A). Platelets in immuno-stained sections of WT clots appeared activated and formed nodes of fibrin bundling and contraction (Figure 2B, left panel). In contrast, platelets in *Rasgrp2*^*-/-*^ clots retained a rounded, unactivated morphology and platelet-fibrin nodules were lacking, even though *Rasgrp2*^*-/-*^ platelets were interacting with fibrin (Figure 2B, middle panel).

**Figure 2:**
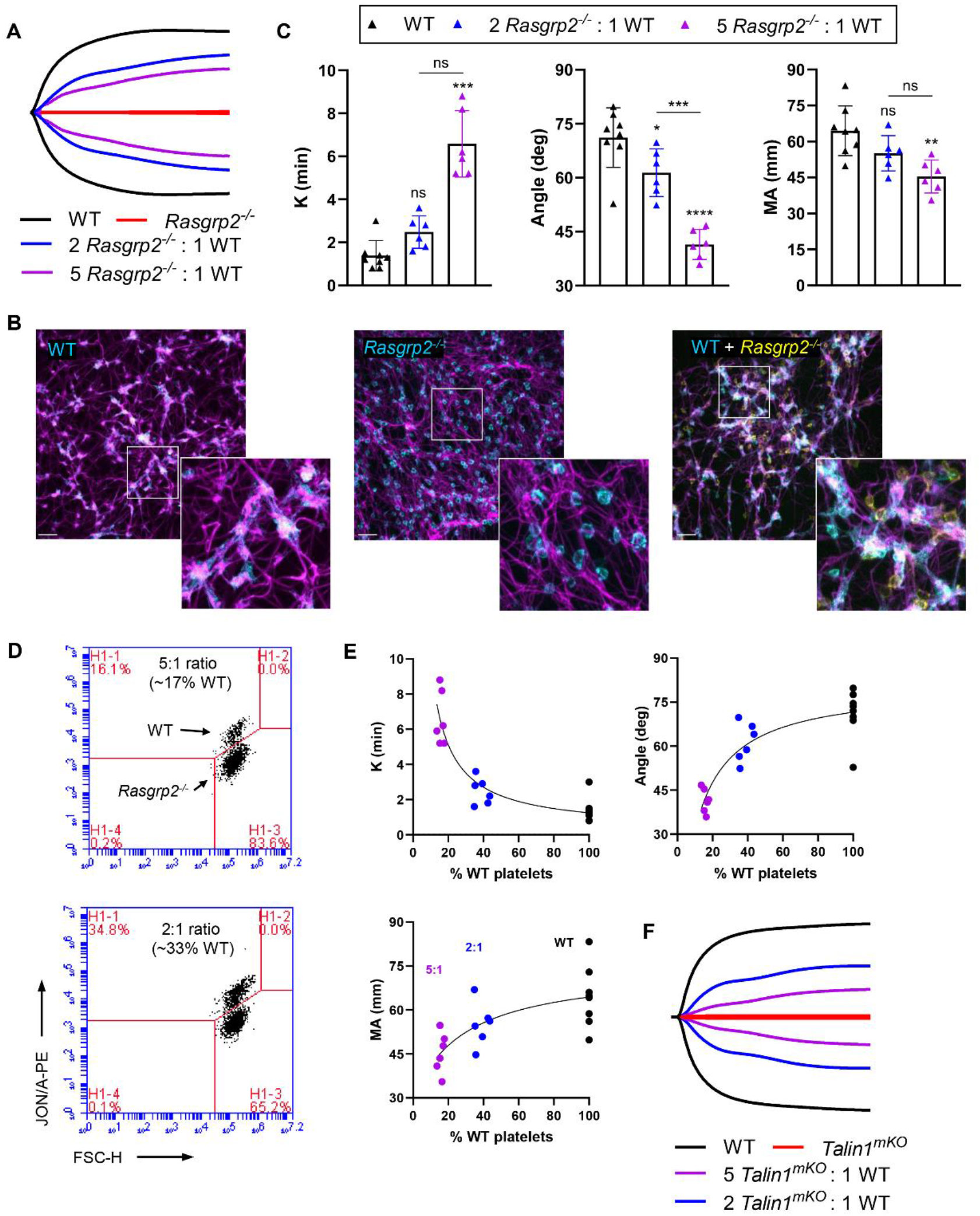
(A) Whole blood (WB) samples from WT and *Rasgrp2*^*-/-*^ mice, kept separate or mixed to achieve specific platelet ratios, were analyzed by TEG-PM with arachidonic acid (AA). Representative TEG-PM traces are shown. (B) Anti-GPIX antibody and AlexaFluor-conjugated fibrinogen were added to WB samples prior to TEG-PM assay, followed by fixation and fluorescence confocal imaging of clots. Platelets are shown in cyan in WT and *Rasgrp2*^*-/-*^ clots, or cyan (WT) and yellow (*Rasgrp2*^*-/-*^) in the mixed sample. Fibrin is shown in magenta in all samples. Scale bars = 10 µm. (C) TEG-PM parameters (K, α-angle, MA) for WT, 2:1 and 5:1 ratios. Parameters for *Rasgrp2*^*-/-*^ samples are not shown due to flatline trace. N = 8 for WT; n=6 for 2:1 and 5:1 ratios; data shown as mean ± SD. Statistical significance was performed by one-way ANOVA with Tukey’s multiple comparisons test for angle and MA, or by Kruskal-Wallis with Dunn’s multiple comparisons test for K. **P<0.01, ***P<0.001, ***P<0.0001, ns-not statistically significant. (D) Platelets were activated with 20 µM ADP in WB in the presence of anti-GPIX (for gating) and JON/A-PE (for activated αIIbβ3) antibodies. Example flow cytometry plots of gated platelets used to determine % WT platelets are shown for 5:1 and 2:1 samples. (E) Plots of TEG-PM parameters versus % WT platelets for individual samples. (F) Representative TEG-PM traces for ratios of WT and *Talin1*^*mKO*^ samples.

To mimic the platelet ratios tested in our previous in vivo study [11], WB samples from *Rasgrp2*^*-/-*^ and WT mice were mixed to obtain ratios of ∼5:1 or ∼2:1 *Rasgrp2*^*-/-*^:WT platelets. At a 2:1 ratio, representing 2/3 *Rasgrp2*^*-/-*^ platelets and 1/3 WT platelets, K and MA improved nearly to WT values, while α remained slightly but significantly reduced compared to WT samples (Figure 2A,C). Consistently, nodules of activated WT platelets and contracted fibrin were observed in clot sections, and rounded, unactivated *Rasgrp2*^*-/-*^ platelets seemed to interact with both WT platelets and fibrin without affecting nodule formation (Figure 2B, right panel). At a 5:1 ratio, K was significantly prolonged compared to WT, although not to a significant extent compared to 2:1 ratio samples (Figure 2C). α remained low and was significantly reduced compared to WT and 2:1 ratio samples. MA improved from <10 for *Rasgpr2*^*-/-*^ to levels comparable to those obtained for 2:1 ratios, though still significantly lower than WT (Figure 2C). To determine the percentage of WT platelets in each sample, mixed WB samples were stimulated with ADP and αIIbβ3 integrin activation was determined by flow cytometry. *Rasgrp2*^*-/-*^ platelets do not activate αIIbβ3 in response to ADP stimulation and thus can be discriminated from WT cells (Figure 2D). When analyzing each TEG parameter against the percentage of WT platelets in each individual sample, the strongest association was observed for K and α (R^2^ = 0.82 and 0.78, respectively), with less robust association for MA (R^2^ = 0.50) (Figure 2E). The same ratios were also established with blood from mice lacking Talin1 in megakaryocytes/platelets (*Talin1*^*mKO*^), which are unable to activate αIIbβ and participate in aggregation or clot contraction (considered GT-like)[13] TEG-PM traces for *Talin1*^*mKO*^:WT ratio samples were comparable to those of *Rasgrp2*^*-/-*^:WT platelets (Figure 2F).

This proof-of-concept study demonstrates that TEG-PM may be a useful method to monitor platelet transfusion ratios and hemostatic efficacy in certain inherited platelet disorders. Several previous clinical reports have described ineffective hemostatic control with platelet transfusion in patients with Glanzmann’s thrombasthenia (GT) or pathological *RASGRP2* variants [6,7,9]. However, most studies do not report the use of any post-transfusion assessment of platelet number or function. A notable exception is a pediatric GT patient undergoing oral surgery, where transfusion of one apheresis unit resulted in a 5:1 GT:normal platelet ratio based on post-transfusion counts. Correction of hemostasis, however, required transfusion of an additional 4 units and a 1:1 ratio [9]. Ex vivo mixing studies suggested that aggregometry may be useful in predicting hemostatic potential in mixed platelet samples, but our studies with murine platelets did not support this conclusion. For some GT and Bernard Soulier Syndrome (BSS) patients, i.e. patients with altered expression of abundant platelet surface glycoproteins, flow cytometry was shown to be a powerful method to monitor transfusion ratios [15,16]. However, more complex analysis platforms would be required to monitor platelet function in the case of mutations to genes coding for intracellular proteins. Another method used to monitor platelet function in the clinic is thromboelastography/elastometry (TEG/ROTEM). While these assays are widely used to monitor platelet function in the context of trauma, a few reports suggest that it can be used to monitor hemostatic potential in GT patients that received platelet transfusion [15,17,18]. In this case, standard TEG or INTEM/EXTEM proved useful as MA/MCF parameters depend on integrin-mediated platelet contraction [19]. However, whether viscoelastic parameters following transfusion correlate with hemostasis in these patients has not been determined. In an adult GT patient undergoing surgery, EXTEM parameters were markedly improved following transfusion of 2 platelet units prior to surgery, but under these conditions the patient suffered profuse bleeding [20]. We recently confirmed integrin-dependence of MA in standard TEG for mouse platelets; however, these studies also showed that standard TEG traces are not affected for *Rasgrp2*^*-/-*^ mice [14], a finding that is consistent with *Rasgrp2*^*-/-*^ *or RASGRP2* mutant platelets having near-normal responses to high concentrations of thrombin and normal contraction [4]. TEG-PM specifically measures the ability of platelets activated by non-thrombin agonists (AA or ADP) to engage and contract reptilase-generated fibrin [21]. Our studies demonstrate that TEG-PM is a powerful method to predict the hemostatic potential in blood samples containing both dysfunctional and healthy platelets, at least in mice. Our results here suggest that K and α were most predictive of a hemostatic platelet ratio. Clinically, TEG-PM is predominantly used to monitor the effects of antiplatelet drugs and the MA value, representing the difference between MA following activation with ADP or AA compared to MA in standard (kaolin) TEG [21], provides a measure of platelet inhibition. We propose that K and α in TEG-PM are better suited than MA to monitor the efficacy of platelet transfusion, as they provide information on the kinetics of platelet-mediated clot contraction, a process that seems highly dependent on the ratio of dysfunctional to healthy platelets. A major barrier could be prevalence of viscoelastic assays in platelet labs; results of a 2014 ISTH survey revealed that while nearly 60% of labs used flow cytometry, <5% reported having a TEG machine [22], although availability would be higher in surgical settings. Future studies will be required to validate our findings here with human samples.

## Author Contributions

R.H. Lee. performed experiments, analyzed data and drafted the manuscript. T. Rudran performed experiments. W. Bergmeier designed studies and wrote the manuscript. All authors edited and approved the manuscript.

## Acknowledgements

We thank Dr. Alisa Wolberg (UNC) for use of the TEG 5000 analyzers. This work was supported by National Institutes of Health (R35HL144976 to WB). All authors declare no conflicts of interest.

